# Correlating Deep Learning-Based Automated Reference Kidney Histomorphometry with Patient Demographics and Creatinine

**DOI:** 10.1101/2023.05.18.541348

**Authors:** Brandon Ginley, Nicholas Lucarelli, Jarcy Zee, Sanjay Jain, Seung Sook Han, Luis Rodrigues, Tezcan Ozrazgat-Baslanti, Michelle L. Wong, Girish Nadkarni, Kuang-Yu Jen, Pinaki Sarder

## Abstract

**Background:** Reference histomorphometric data of healthy human kidneys are largely lacking due to laborious quantitation requirements. Correlating histomorphometric features with clinical parameters through machine learning approaches can provide valuable information about natural population variance. To this end, we leveraged deep learning, computational image analysis, and feature analysis to investigate the relationship of histomorphometry with patient age, sex, and serum creatinine (SCr) in a multinational set of reference kidney tissue sections.

**Methods:** A panoptic segmentation neural network was developed and used to segment viable and sclerotic glomeruli, cortical and medullary interstitia, tubules, and arteries/arterioles in the digitized images of 79 periodic acid-Schiff-stained human nephrectomy sections showing minimal pathologic changes. Simple morphometrics (e.g., area, radius, density) were quantified from the segmented classes. Regression analysis aided in determining the relationship of histomorphometric parameters with age, sex, and SCr.

**Results:** Our deep-learning model achieved high segmentation performance for all test compartments. The size and density of nephrons and arteries/arterioles varied significantly among healthy humans, with potentially large differences between geographically diverse patients. Nephron size was significantly dependent on SCr. Slight, albeit significant, differences in renal vasculature were observed between sexes. Glomerulosclerosis percentage increased, and cortical density of arteries/arterioles decreased, as a function of age.

**Conclusions:** Using deep learning, we automated precise measurements of kidney histomorphometric features. In the reference kidney tissue, several histomorphometric features demonstrated significant correlation to patient demographics and SCr. Deep learning tools can increase the efficiency and rigor of histomorphometric analysis.

**Significance statement:** Although the importance of kidney morphometry is well explored in disease contexts, the definition of variance in reference tissue is not. Advancements in digital and computational pathology have rendered quantitative analysis of unprecedented tissue volumes via the single press of a button. The authors leverage the unique benefits of panoptic segmentation to perform the largest ever quantitation of reference kidney morphometry. Regression analysis identified several kidney morphometric features that varied significantly with patient age and sex, and the results suggested that the set size of nephrons might depend more intricately on creatinine than previously thought.

## Introduction

Diagnostic renal pathology relies on recognizing histologic findings that deviate from the expected range for “normal” or “healthy” tissue, hereby defined as reference tissue with none to minimal histopathologic abnormality. However, detailed reference histomorphometric data of healthy human kidneys are largely lacking due to laborious quantitation requirements. With recent advances in the technical capabilities and performance of deep learning (DL)-based image analysis, computational pathology has emerged as a potential feasible and scalable method to automate histomorphometric quantitation and provide accurate reference kidney histomorphometric data. If applied to a large and diverse sample of healthy human kidneys, this approach can provide insights into histomorphometric natural variance within populations and subgroups, which may correlate to clinical parameters and disease susceptibility. Similar detailed quantitation in disease states may provide additional diagnostic and prognostic information that are not currently available or feasible for routine diagnostic renal pathology practice. However, the utility of such data can only be recognized if reliable reference values are available for comparison.

In this proof-of-concept study, we developed a DL-based image analysis model for conducting comprehensive segmentation of renal histomorphometry and applied it to whole slide images (WSIs) of reference kidney tissue in order to automate the measurement of simple histomorphometric features. These quantified features were then correlated to patient demographics and serum creatinine values to assess their biological relevance.

## 1. Methods

This study was approved by the institutional review board at the University of Florida with waiver of informed consent.

### 1.1 Kidney tissue sections

All deidentified kidney tissue sections in this study were obtained from the pathology archives of the University of California, Davis, Centro Hospitalar e Universitário de Coimbra, and Seoul National University Hospital. The original tissue was formalin-fixed, paraffin-embedded (FFPE), and sectioned at 2-4 μm in thickness. The cases and stains used are detailed below.

### 1.2 Image data

WSIs were generated by scanning the glass slides with a whole slide brightfield microscopy image scanner (Aperio, Leica, CA). Spatial annotation of tissue sections was performed in Aperio^®^ImageScope and saved as ImageScope compatible XML files.

#### 1.2.1 Training data

For segmentation training, 190 WSIs were collected, including 53 diabetic nephropathy, 39 lupus nephritis, and 11 transplant surveillance needle core biopsies (total tissue area, 1100 mm^2^), 58 large sub-WSIs manually cropped from healthy portions of 33 reference kidneys (total tissue area 369 mm^2^), 23 small sub-WSIs of 5 H&E needle core biopsies (total tissue area 34 mm^2^), 2 small sub-WSIs from a silver-stained biopsy (2 mm^2^), and 4 sub-WSIs from a trichrome biopsy (1.4 mm^2^). These slides were annotated in their entirety for viable and sclerotic glomeruli, cortical and medullary interstitia, tubules, and arteries/arterioles. Overall, 1506 mm^2^ of kidney tissue was annotated.

#### 1.2.2 Performance analysis and ground-truth data

Ten PAS transplant surveillance kidney biopsy sections (9 needle cores, 1 wedge) were used for performance measurement. Five patients had serum creatinine at the time of biopsy of >2 mg/dL, and 5 patients were selected to have “histologically normal” biopsy findings as per the original case report in order to measure performance both in diseased and healthy states. None of these slides were included in training the algorithm. Glomeruli <1500 μm^2^ in the area and glomerular fragments that were not physically contiguous with the biopsy core were not considered. Very small diameter arterioles were difficult to discern from capillaries; therefore, the smallest vessels annotated as arterioles for the purpose of performance evaluation were those >200 μm^2^ in area and displaying at least one full circumferential layer of smooth muscle cells, barring the loss of smooth muscle to disease. The smallest annotatable tubule was also defined as those >200 μm^2^ in area, as below this threshold the objects could not be confidently determined as tubules. The area thresholds discussed herein were applied to the neural network segmentation outputs to filter the structures for performance evaluation.

#### 1.2.3 Reference kidney data

The reference kidney tissue sections consisted of archived glass slides of the renal parenchyma uninvolved and away from the renal tumor of human tumor nephrectomy specimens. These cases were screened to have no evidence of hydronephrosis, infectious disease, or proteinuria. The renal pathologist screened the slides to include only cases with minimal pathologic changes (e.g., no tumor, no significant preservation or processing artifact, and <5% interstitial fibrosis and tubular atrophy [IFTA]). In total, reference sections from 79 unique subjects were included, with a tissue area totaling 17,208 mm^2^.

### 1.3 Renal multicompartment segmentation

#### 1.3.1 Tissue detection

The renal tissue regions were detected from the background by creating a low-resolution thumbnail (16x downsample) of the entire WSI and transforming it to the hue, saturation, and value color space.^1^ From the resultant image, the total tissue area was measured by thresholding the saturation channel at 0.05, summing all of the pixels in the resultant binary mask, and converting the output to mm^2^. To identify image regions of tissue for deep learning processing, the saturation channel was further blurred with a Gaussian filter (*σ* = 5) to create a loose buffer zone around the tissue edge. The blurred saturation image was converted to a binary mask by thresholding at 0.05. This low-resolution mask was gridded into a set of tiles based on the desired training or testing patch size for renal multicompartment segmentation, amount of overlap between patches, and tolerable percent of non-tissue per patch. For training, image size was specified as 1200 × 1200 pixels and an overlap of 50% between adjacent tiles was allowed. For testing, image size was specified as 2048 × 2048 pixels, with 10% overlap between patches. For both training and testing, any tile with greater than 99% background was excluded from further processing.

#### 1.3.2 Data loading for DL

Network training was orchestrated using the Dectron2 library for PyTorch,^2^ which implements convenient functions for training and evaluating a panoptic feature pyramid architecture. A custom dataloader to extract image crops and associated labels from WSIs and XMLs was designed to feed network training *‘on the fly’* rather than saving image crops to disk, resulting in reduced memory overhead and disk usage, as well as allowing the added convenience of implementing balanced data sampling routines both at the whole-slide and target-class levels.

Given that the cross-sectional area of kidney tissue sections consists of mostly tubules, a data selection routine for class balance was required to prevent an underfit classifier on non-tubule targets and performed as follows: For each patch requested during training, one slide from the training set was selected randomly. Next, with a 50% probability, either a random slide tile was extracted from the tissue area or the tile was selected to be centered on a randomly selected artery/arteriole or glomerulus. All random sampling was performed using a uniform distribution.

#### 1.3.3 Training

Network weights were initialized to a model pretrained on the COCO dataset^3^ available in the Detectron2 library, which had a ResNet-50 backbone and was originally trained with the 3X learning rate schedule. The following modifications were made to the network architecture, which differed from the stock configuration of Detectron2: the anchor generator sizes were specified as 32, 64, 128, 256, 512, and 1024; the respective region proposal network’s input layers for these anchors were specified as p2, p3, p4, p5, p6, and p6; the anchor generator aspect ratios were specified as 0.1, 0.2, 0.33, 0.5, 1, 2, 3, 5, 10; and the anchor generator angles were specified as −90, −60, −30, 0, 30, 60, and 90. No image resizing was performed, and training was performed with batch size four and region of interest head batch size 64. Several image augmentations were performed to improve network robustness to unseen test variations (further discussed in the ***Supplementary Document***). A similar training schedule was followed as was laid out in the original implementation^4^ for training on the COCO dataset. Starting from the COCO pretrained model, the network was trained for a total of 350 thousand steps, with a step learning rate policy starting at 0.0025 and dropping by one-tenth upon reaching 100 thousand, 200 thousand, and 300 thousand steps. Glomeruli (viable and sclerotic), tubules, and arteries/arterioles were specified as instance-type segmentation objects, and the interstitium and slide background were specified as semantic-type segmentation objects.

#### 1.3.4 Testing

The custom data loader described in ***Section 1.3.2*** was repurposed for prediction on test biopsy data by converting its output to yield each tile in a WSI grid once. Tiles were sent to the trained deep learning network for prediction, and the corresponding predictions filled into a high-resolution segmentation mask within the WSI. Predictions in overlapped regions of tiles were resolved by clipping the trailing and leading edges of overlap halfway. Additionally, the panoptic network’s region of interest head (see ^2^ and ^4^ for further details) threshold was set to 0.01 to maximize the number of detected instances. All objects in the final high-resolution mask were converted to their corresponding boundary contour vertices and stored in an XML file compatible with Aperio ImageScope (Leica Biosystems, Nussloch, Germany) or in JSON files compatible with HistomicsUI.^5^

### 1.4 Segmentation performance analysis

Multicompartment segmentation performance was assessed both pixel-wise and instance-wise for a comprehensive performance evaluation of our deep learning pipeline.

#### 1.4.1 Pixel-wise performance analysis

Whole slide manual annotations of the instance segments were compared pixel-wise against network output whole slide predictions for each class using a one-versus-all approach. The true/false positive/negative pixels were pooled across the entire dataset to calculate the final reported performance values, including sensitivity, specificity, precision, negative predictive value, Matthew correlation coefficient and Dice coefficient.^6^

#### 1.4.2 Instance-wise performance

Instance performance calculations were evaluated in the cortex of each WSI only, as medulla is not used routinely for diagnostic purposes. Network-predicted instances were annotated with a dot marker if the prediction was incorrect for any reason. The types of error for each instance prediction were broken down into fused instances, partially detected instances, missed instances, and wrong classifications, and counted at the WSI level. Partial, fused, and false class percentage error rates were calculated as 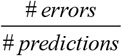. Missed percentage error rates were calculated as 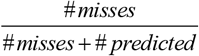. Total error rate was calculated as 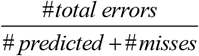, where #*total errors* was defined as the sum of #*errors* and #*misses*.

### 1.5 Reference kidney morphometry

Reference kidney morphometrics were quantified using the saved contour representation of segmented object boundaries for each WSI. The full list of tested features is available in the ***Supplementary Document***. A generic overview of our object feature quantification strategy is discussed below. Note that any glomerular predictions contained within medulla were algorithmically eliminated.

#### 1.5.1 Object diameter

Calculation of diameters for segmented spherical objects (i.e. glomeruli) is straightforward. Diameter measurements of non-spheroid objects (e.g., tubules and vessels) are complex and subject to bias. Thus, an automated method that can measure object diameters and minimize sampling bias but maximize application to various sectioned orientations of histologic structures was developed, reliant on a morphometric processing method called distance transform.^7, 8^ The distance transform takes each object pixel and measures the distance to the closest boundary point. The result at every pixel describes the largest radius of a circle centered at that pixel and inscribed within the object. The maximum of all of these pixel values is the radius of the largest circle that be inscribed within the object. We defined tubular and vessel diameter using the diameter of the largest circle. The ***Supplementary Document*** discusses examples of the distance transformation for varied tubule segments.

#### 1.5.2 Object area

Segmented object areas were calculated from contour vertices using Green’s theorem.^9^ To compute the cortical interstitial area, the aggregate area of objects contained within the cortex (i.e. glomeruli, tubules, and arteries/arterioles) was subtracted from the total cortical area. Similarly, the medullary interstitial area was calculated by subtracting the aggregate area of medullary tubules from the total medullary area.

#### 1.5.3 Object densities

Enumeration and quantification of segmented objects were normalized to the total tissue area that contained the objects, which represents respective object densities. The simplest of these metrics was the division of the number of counted glomeruli, tubules, or arteries/arterioles, or their summed areas, by the observed area over which they were distributed (either cortical area, medullary area, or both). To compute the interstitial density, the calculated cortical or medullary interstitial areas were divided by the total cortical or medullary contour areas, respectively.

#### 1.5.4 Arterial/arteriolar luminal ratio

Arterial/arteriolar luminal ratio was calculated as the radius of the artery/arteriole lumen divided by the radius of the entire segmented vessel. To identify the luminal area, the corresponding RGB image region for each artery/arteriole segmentation was extracted, transformed to LAB colorspace,^7, 8^ and the lightness channel of the LAB colorspace was thresholded at 70, yielding a segmentation of the white regions in the vessel. Vessels with overall image width or height >5000 pixels were excluded from this analysis due to the network commonly detecting these vessels as fragments, limiting the ability to properly segment lumina.

### 1.6 Statistical analysis

Multivariable linear regression analyses were performed using age, sex, and SCr as predictor variables, institutional source of data as fixed effects, and morphometric measurements as outcome. Standard errors were calculated using a cluster robust method.^10^ All statistical analysis was performed in R.

Statistical significance analysis of the data was shown in ***Table 3*** and ***Supplementary Table 1***. The Fisher’s exact test was used to test independence between categorical variables, while the analysis of variance (ANOVA) was used to test differences across institutions for continuous variables. Bonferroni correction was used to adjust for multiple comparisons.^11^

### 1.7 Hardware and computational time

Computational processing was performed on a Linux distribution (Ubuntu 16.04) computer with an Intel(R) Xeon(R) Silver 4114 CPU with 40 cores at 2.20 GHz, 64 GB of RAM, and 64 GB of swap memory. Network operations were performed on a Geforce RTX 2080 Ti GPU (11 GB memory). Multicompartment segmentation of a typical biopsy section image of size 2 mm^2^ using our pipeline takes 30 min and a typical nephrectomy of size 100 mm^2^ takes 12 hours. Computation of all morphometric data from one section takes roughly between 30 sec and 4 min, heavily depending on the time spent calculating features on tubules, which varies between 7K and 82K in our dataset.

## 1.8 Data availability

All 79 reference WSIs, and segmented renal micro-compartments in XML format are available at https://bit.ly/3YD4r6a.

## 2. Results

### 2.1 Segmentation model performance

To assess the performance of the segmentation model, a holdout test set of 10 PAS-stained human kidney transplant biopsies was used, comprised of 5 cases from patients with >2 mg/dL SCr at the time of biopsy and 5 cases with minimal to no histologic abnormalities as determined by the renal pathologist. This strategy was employed to evaluate the network performance for both normal and diseased states. Examples of the kidney segmentation output in the test biopsies are shown in ***Fig. 1***. To quantitate the model performance, every slide was manually reviewed and all instances of incorrect predictions were tabulated. Possible sources of instance error included incomplete segmentation of the full boundary (partial), complete non-detection (missed), fusion of two boundaries that should be distinct (fused), or correct placement of the boundary with incorrect class assignment (false class). The prediction errors across the 10 slides are detailed in ***Table 1***.

**Fig. 1.**
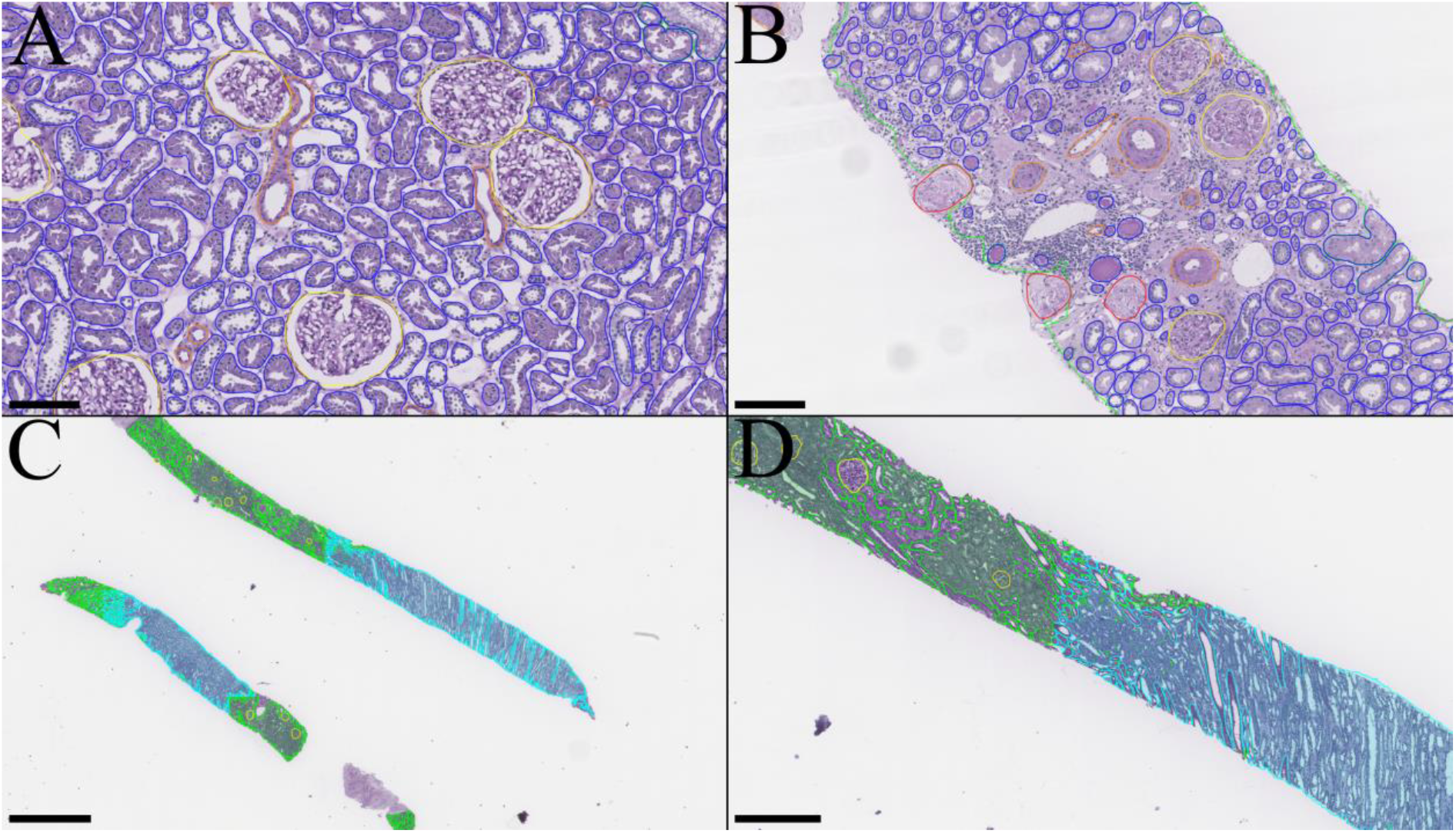
Panoptic segmentation of test set kidney biopsies. A) Instance predictions in a kidney biopsy showing healthy / normal parenchyma. B) Instance predictions in a kidney biopsy from a patient with creatinine > 2mg/dL. C) Low-resolution demonstration of corticomedullary semantic segmentations. D) Zoomed inset from the top core in C. Green: cortex; cyan: medulla; yellow: viable glomerulus; red: sclerotic glomerulus; blue: tubule; orange: artery/arteriole. Scale bars: A, 150μm; B, 150μm; C, 1.5mm; D, 500μm.

**Table 1.**
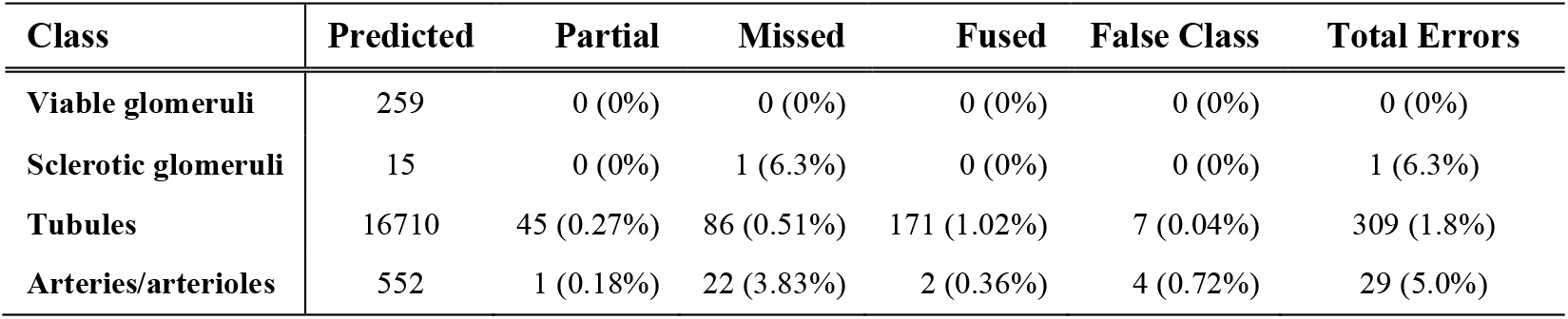
Instance error rates on the test set. Values reported as absolute count (%).

For the viable glomerulus class, the network identified every glomerulus while making no false detections. The network also performed well on tubule segmentation with a 1.8% total error rate. Most tubule segmentation errors were due to the fusion of tubular boundaries, which occurred when 2 or more tubules were in very close proximity, often showing essentially no appreciable intervening interstitium (***Fig. 2A***). An appreciable minority of tubule segmentation errors were missed tubules; however, these tended to be very small atrophic tubules, very small caliber tubules in the medulla, or extremely tangentially sectioned tubules (***Fig. 2B***). Errors in the segmentation of arteries/arterioles mainly consisted of missed instances, typically of very small vessels that were bordering on the size of capillaries (***Fig. 2C***).

**Fig. 2.**
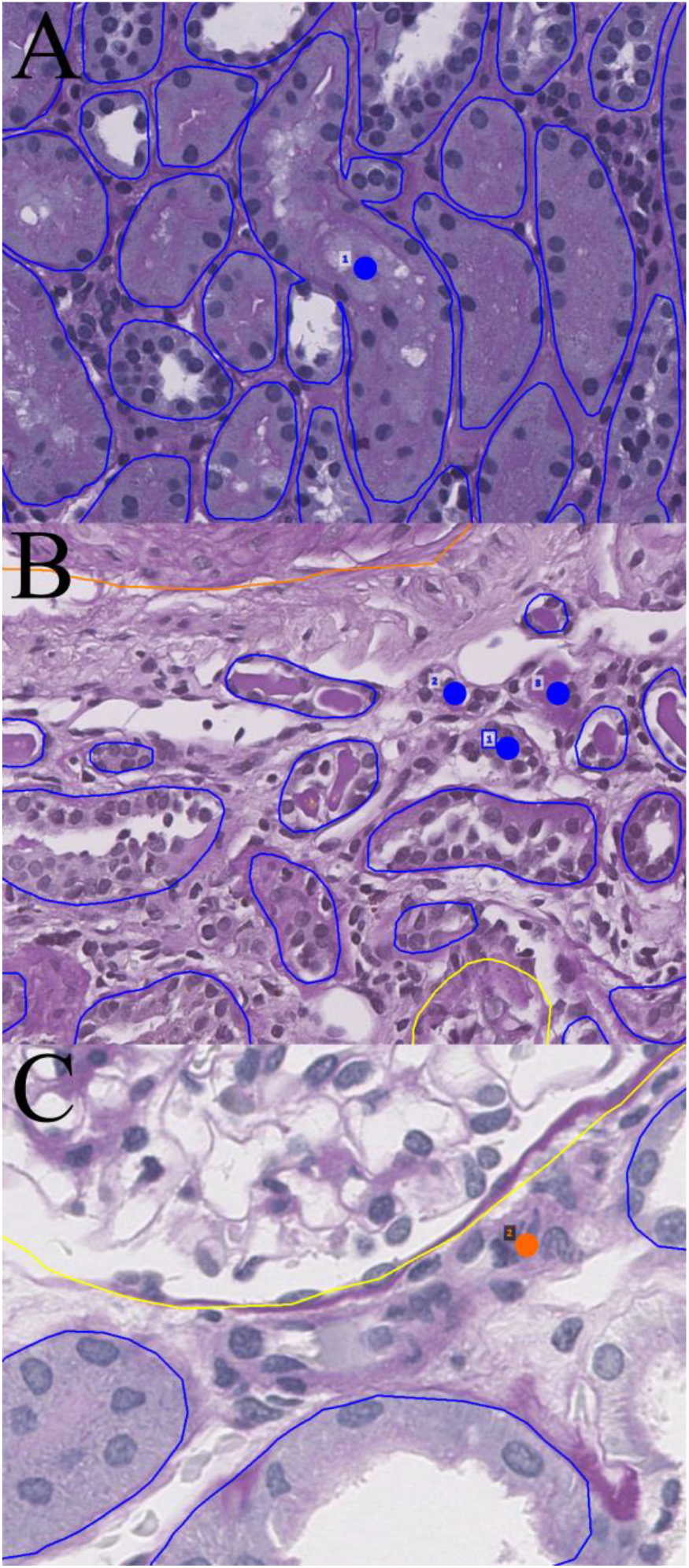
Network errors. Dots indicate missed structure. A) Network fusion of tubular boundaries when basement membranes abut and morphologies are grossly dissimilar. B) Network misses on small atrophic tubules. C) Network miss on small arteriole bordering capillary size. Blue: tubule; orange: artery/arterioles.

The network was least performant in detecting sclerotic glomeruli, although the error rate is misleading since only one sclerotic glomerulus was missed of a total 16 present in the 10 test cases. The one missed instance was a sclerotic glomerulus cut in half at the biopsy edge.

After the detection of erroneous instances, their boundaries were manually corrected to measure a pixel-by-pixel performance of the segmentation output. These values are reported in ***Table 2*** and essentially reflect the instance error rate results.

**Table 2.** Pixel-wise performance metrics compared against renal pathologist.

| **Class** | **Sensitivity** | **Specificity** | **Precision** | **NPV** | **MCC** | **Dice** |
| --- | --- | --- | --- | --- | --- | --- |
| **Glomeruli** | 0.998 | 1 | 1 | 1 | 0.999 | 0.999 |
| **Sclerotic glomeruli** | 0.948 | 1 | 0.996 | 1 | 0.972 | 0.972 |
| **Tubules** | 0.995 | 1 | 0.999 | 1 | 0.997 | 0.997 |
| **Arteries/arterioles** | 0.984 | 1 | 0.994 | 1 | 0.989 | 0.989 |

**Abbreviations:** NPV: Negative predictive value, MCC: Matthew's correlation coefficient, Dice: Dice coefficient (F1-score).

### 2.2 Reference kidney morphometrics

Using the panoptic segmentation model, measurement of histomorphometric parameters was performed for a set of reference kidneys. Since kidney tissue from individuals with no renal disease is typically not available, this study was performed on sections of renal parenchyma uninvolved and away from the renal tumor of human tumor nephrectomy specimens. Inclusion and exclusion criteria of the reference kidney are detailed in ***Section 1.2.3*** and were designed to minimize the presence of abnormal histologic findings. In total, 79 multinational nephrectomy cases were included, derived from three international institutions, and each kidney section was stained with PAS. Quantified features for the reference kidney cases are tabulated in ***Table 3***. Examples of whole slide segmentation for reference kidneys are shown in ***Fig. 3***.

**Table 3.**
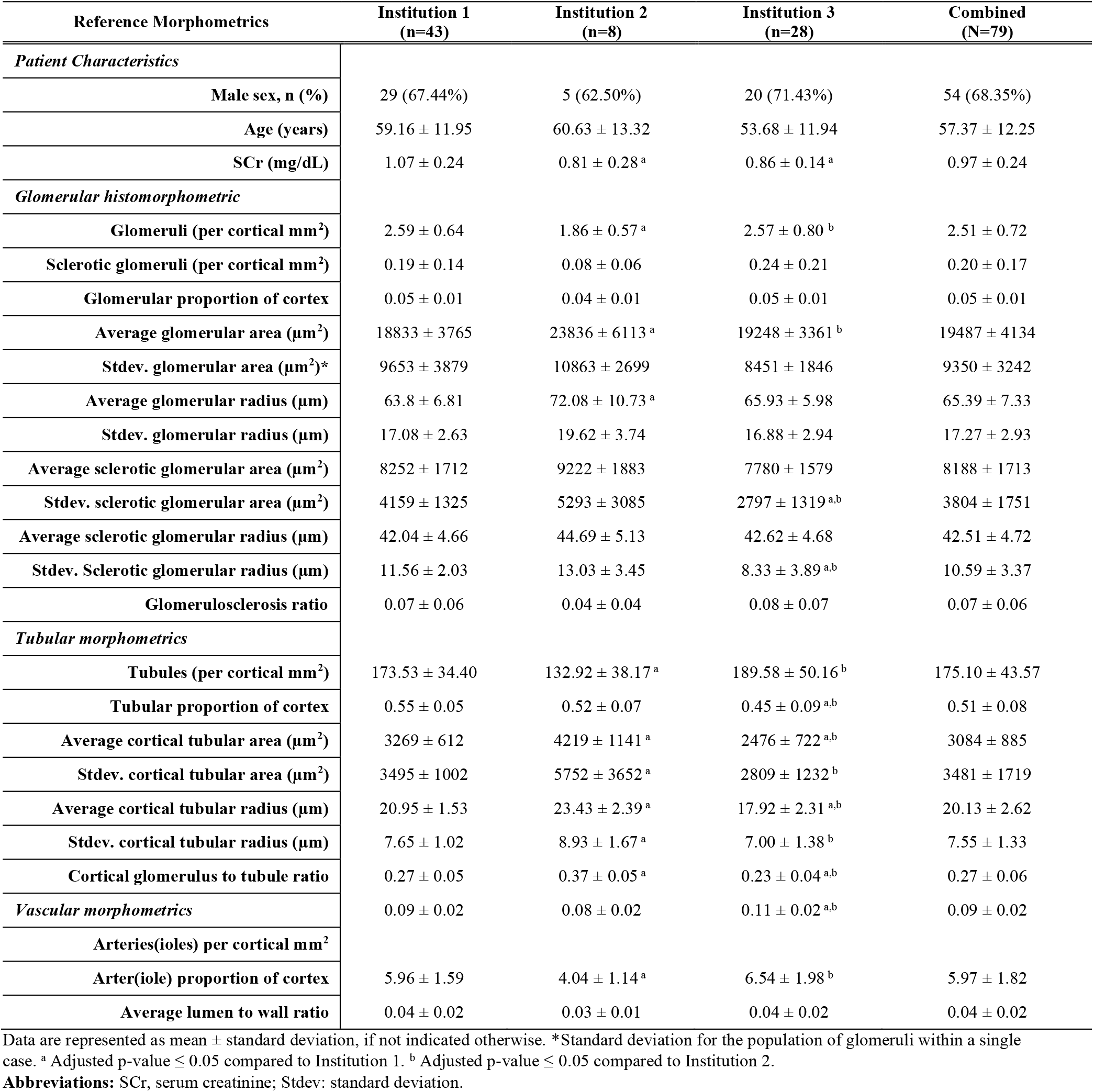
Reference morphometrics (*n* = 79 subjects; one whole slide image per subject).

**Fig. 3.**
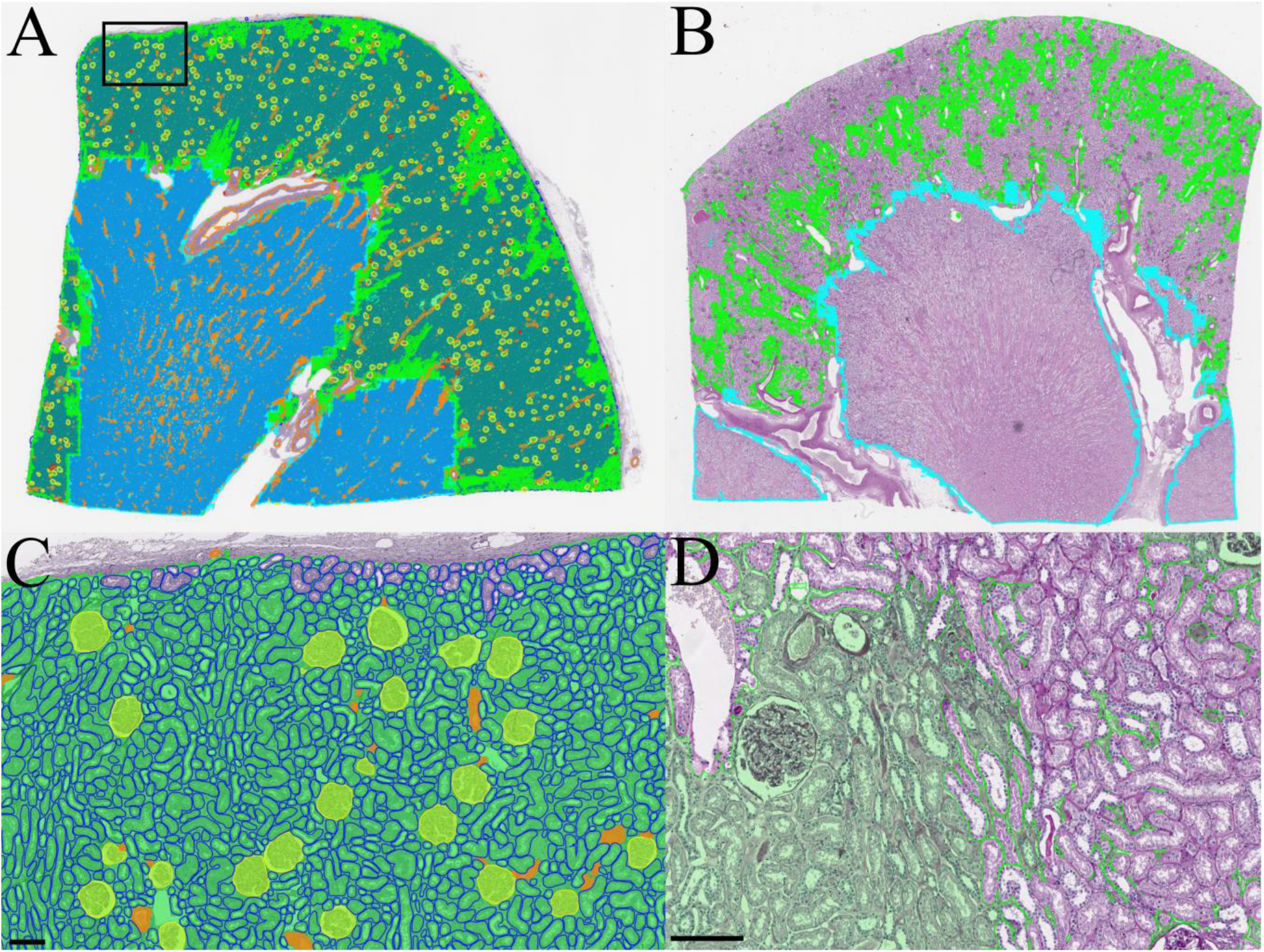
Whole section segmentations for PAS-stained kidney nephrectomies. A) Thumbnail of whole segmentation mask for a reference kidney. Tubules are rendered in the background to prevent them from overwhelming the visibility of other structures. B) Thumbnail of patchy interstitial segmentation in a kidney with many tubules flush back to back. C) Zoomed region from A showing segmentation of viable glomeruli, tubules, arterioles, and cortical interstitium. D) Zoomed region from B showing interstitium at left fused by contour retrieval after tile stitching process, where interstitium at right is patchy due to flushly abutting tubules. Green: cortical interstitium; cyan: medullary interstitium; yellow: viable glomerulus; red: sclerotic glomerulus; blue: tubule; and orange: artery/arteriole. Scale bar, 150μm.

All three institutional patient cohorts displayed similar proportions of males versus females and age distributions. Serum creatinine values, albeit varied between the institutions, were measured to be in normal range. For the histomorphometric parameters, the vast majority were similar between institutions, with a few notable exceptions discussed below.

For glomeruli, the average number of glomeruli per mm^2^ of cortical tissue was 2.5, of which the average number of sclerotic glomeruli per mm^2^ was 0.2. These values equate to a glomerulosclerosis rate of 7.3%, which matches expectations for an age range of 50 to 60 years old.^12^ Glomerular area and radius ranged between ∼18800 μm^2^ and 23800 μm^2^ and 64 μm and 72 μm, respectively. Of note, the glomerular density varied slightly when comparing institutions in a pattern that was inversely proportional to the measured average glomerular size.

For tubules, the average number per cortical mm^2^ also varied significantly across institutions, ranging from ∼130 to ∼190, again being inversely proportional with the measured average tubule size. The average area and radii of tubules ranged from 2476 μm^2^ to 4219 μm^2^ and 17.92 μm to 23.43 μm, respectively.

Similarly, the number of observed arteries and arterioles per mm^2^ ranged from 4 to 6, and is found to be inversely proportional with average nephron size. That is, kidneys with larger nephrons have lower densities of arterioles and arterioles. However, we also found those arteries and arterioles to have proportionally much wider lumens, as the ratio of luminal width to overall vessel width was higher, namely, 0.37 as quantified for *Institution 2* vs 0.23 and 0.27 for *Institution 1* and *3*, respectively.

### 2.3 Histomorphometric variation across patient demography and serum creatinine

We next used a series of adjusted linear regressions to determine if histomorphometric measurements made on our reference kidney cohort correlate with basic patient information. This part of the study incorporated patient age, sex, and serum creatinine as input variables, morphometric measurements as output values, and institutional data source as fixed effects. ***Table 4*** summarizes model parameters for this regression analysis.

**Table 4.** Correlation of reference morphometrics to age, sex, and serum creatinine. Women were coded with 0 and men with 1 in this study.

| Outcome ($R^2$) | Age $\beta$ (95% CI) \| $p$ | Sex $\beta$ (95% CI) \| $p$ | SCr $\beta$ (95% CI) \| $p$ |
| --- | --- | --- | --- |
| *Glomerular histomorphometric* |  |  |  |
| Mean glomerular area ($\mu\text{m}^2$) (0.31) | 14.2 (-164.3, 192.6) \| 0.765 | 1660.1 (-306.8, 3627.1)† \| 0.068 | **5874.7 (2457.2, 9292.3)\* \| 0.018** |
| Mean glomerular radius ($\mu\text{m}$) (0.31) | 0.02 (-0.32, 0.36) \| 0.826 | 3.02 (-0.92, 6.97)† \| 0.081 | **10.82 (3.64, 18)\* \| 0.023** |
| Stdev. glomerular area ($\mu\text{m}^2$) (0.09) | 17.6 (-52, 87.2) \| 0.391 | **1384.9 (672.4, 2097.5)\* \| 0.014** | **-1106 (-1770.4, -441.5)\* \| 0.019** |
| Glomerular area density (0.05) | 0 (-0.0002, 0.0002) \| 0.877 | 0.0015 (-0.0066, 0.0097) \| 0.504 | **-0.007 (-0.0104, -0.0036)\* \| 0.012** |
| # Glomeruli per cortical $\text{cm}^2$ (0.21) | 0 (-0.004, 0.004) \| 0.984 | -0.059 (-0.213, 0.094) \| 0.237 | **-0.371 (-0.648, -0.095)\* \| 0.029** |
| # Sclerotic glomeruli per cortical $\text{cm}^2$ (0.23) | 0.0022 (-0.0019, 0.0064) \| 0.146 | 0.009 (-0.0011, 0.0191)† \| 0.061 | 0.0043 (-0.2268, 0.2354) \| 0.943 |
| Mean sclerosed glomerular radius ($\mu\text{m}$) (0.05) | 0.052 (-0.203, 0.308) \| 0.472 | 1.284 (-5.013, 7.581) \| 0.473 | **2.95 (1.899, 4.002)\* \| 0.007** |
| Glomerulosclerosis ratio (0.18) | 0.0018 (-0.0006, 0.0041)† \| 0.082 | **0.0134 (0.0021, 0.0246)\* \| 0.036** | 0.0339 (-0.1596, 0.2275) \| 0.529 |
| *Tubular morphometrics* |  |  |  |
| Cortical tubular area density (0.34) | 0.0003 (-0.0051, 0.0057) \| 0.827 | -0.007 (-0.0806, 0.0665) \| 0.721 | 0.0813 (-0.0286, 0.1912)† \| 0.086 |
| Mean cortical tubular area ($\mu\text{m}^2$) (0.43) | -5.84 (-59.81, 48.13) \| 0.687 | 11.24 (-1199.2, 1221.67) \| 0.972 | 1020.72 (-156.15, 2197.59)† \| 0.065 |
| Mean cortical tubular radius ($\mu\text{m}$) (0.55) | 0.007 (-0.152, 0.165) \| 0.876 | 0.289 (-2.991, 3.569) \| 0.741 | **2.983 (2.197, 3.77)\*\* \| 0.004** |
| Stdev. cortical tubular radius ($\mu\text{m}$) (0.26) | | | |
| Mean medullary tubular radius ($\mu\text{m}$) (0.61) | -0.001 (-0.031, 0.03) \| 0.924 | 0.04 (-0.657, 0.738) \| 0.826 | **1.632 (0.014, 3.251)\* \| 0.049** |
| Stdev. medullary tubular area ($\mu\text{m}^2$) (0.22) | -10 (-27.4, 7.5) \| 0.133 | 512.3 (-232.4, 1257)† \| 0.098 | -436.5 (-2161.4, 1288.4) \| 0.39 |
| Mean arterial(olar) lumen to wall ratio (0.42) | -0.0006 (-0.0045, 0.0034) \| 0.604 | -0.0261 (-0.0528, 0.0005)† \| 0.052 | 0.0178 (-0.132, 0.1676) \| 0.66 |
| *Vascular morphometrics* |  |  |  |
| Glomerulus to cortical tubule area ratio (0.31) | 0 (-0.0006, 0.0005) \| 0.754 | 0.0038 (-0.0068, 0.0145) \| 0.261 | **-0.0245 (-0.0398, -0.0093)\* \| 0.02** |
| # Artery(ioles) per cortical $\text{cm}^2$ (0.18) | 0.005 (-0.001, 0.012)† \| 0.062 | **0.169 (0.005, 0.333)\* \| 0.047** | -0.557 (-1.864, 0.75) \| 0.208 |
| Cortical artery(iole) area density (0.05) | **0.0002 (0.0001, 0.0003)\* \| 0.009** | 0.0018 (-0.0178, 0.0213) \| 0.736 | -0.0011 (-0.0209, 0.0187) \| 0.834 |

Symbols/abbreviations: Stdev., standard deviation; \*\*significance <0.005, \* significance <0.05, † significance < 0.1, $\beta$ : regression coefficient, CI: confidence interval.

Several kidney histomorphometric parameters, especially those related to glomerular and tubular size as well as glomerular density, were significantly associated with serum creatinine. For instance, patients with lower serum creatinine (presumably better renal function) tended to have smaller glomeruli (i.e., smaller glomerular area and radii) but higher numbers of glomeruli per renal cortical area (i.e., high glomerular density). Similarly, tubular radii were larger in patients with higher serum creatinine levels. Interestingly, when looking at the standard deviation for the distribution of glomerular and tubular sizes within a kidney tissue section, glomerular size distribution varied less when patients had high serum creatinine levels while tubular size distribution varied more with higher serum creatinine levels.

Fewer significant associations were seen for patient sex and age. The glomerulosclerosis ratio (essentially the percentage of glomerulosclerosis) as well as the density of arteries/arterioles were significantly higher for men than women. Also, the standard deviation for the glomerular area of any given patient tended to be higher for men than women. In terms of age, the only parameter that showed significant association was cortical arterial/arteriolar area density, which was positively correlated to age, meaning that with an increase in age the total area of arteries and arterioles occupying a given amount of cortex increases. The percentage of glomerulosclerosis showed a positive trend with age, although this relationship was not statistically significant.

The distribution of histomorphometric parameters within each patient’s kidney tissue section was also found to have an increasing trend, which is likely due to arterioles thickening with age causing them to be more prominently detected by the segmentation algorithm kernel density estimations coded by color reflecting the patient’s serum creatinine (***Fig. 4***). As illustrated in ***Fig. 4A***, the average and spread (i.e. standard deviation) of cortical tubular radii were both higher in patients with higher serum creatinine values. Similarly, the average glomerular radii (***Fig. 4B***) were higher in patients with higher serum creatinine, but the standard deviations were slightly lower in patients with higher SCr. This observation suggests that as glomeruli hypertrophy to compensate for increased creatinine, they may reach an expansion limit of ∼100 μm in radius. Interestingly, the average sclerotic glomerulus radius was also dependent on creatinine.

**Fig. 4.**
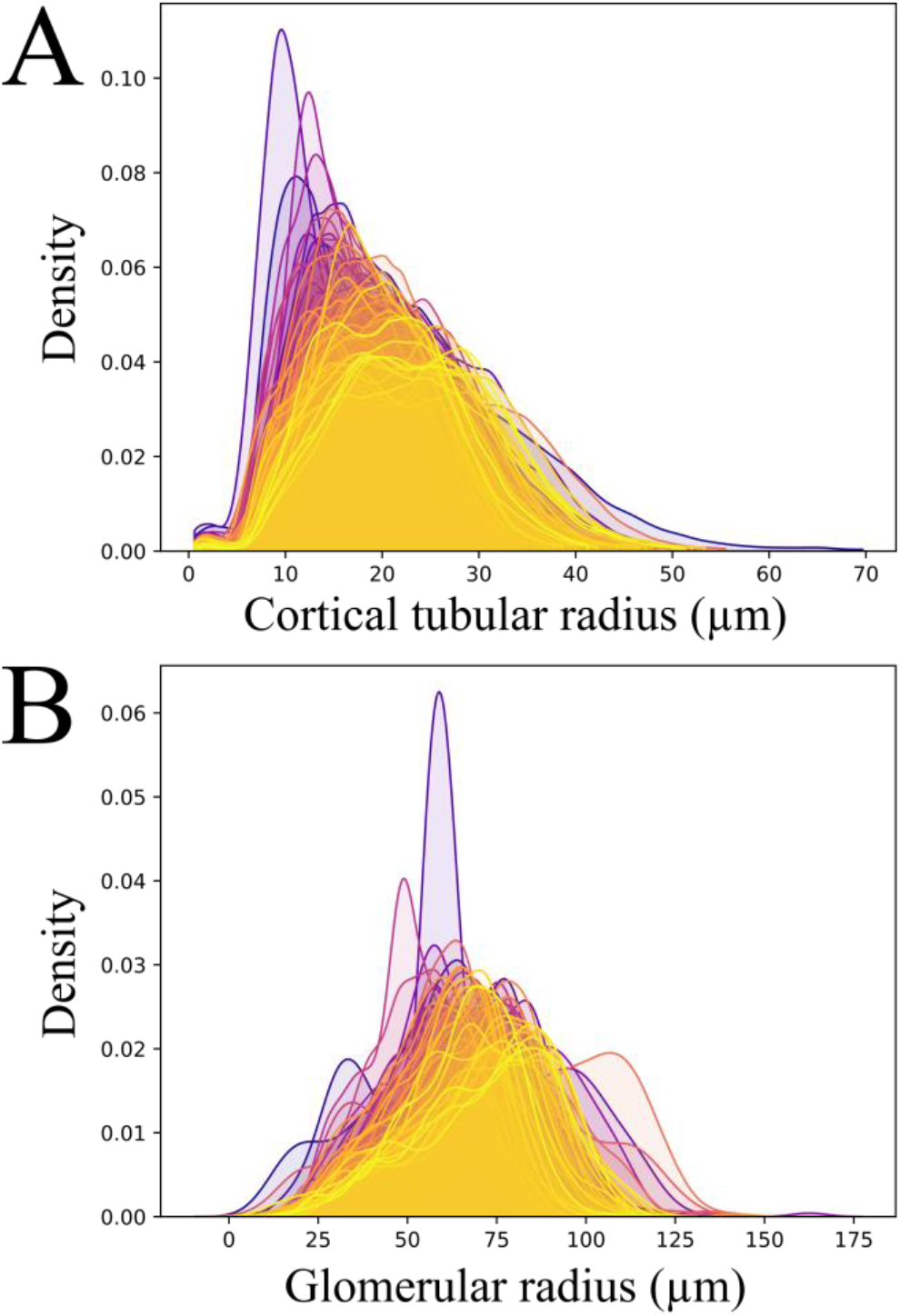
Probability density estimates for nephron radius per patient. Color is coded by creatinine with the lowest values in blue and the highest values in yellow as indicated by the bottom legend. A) Radii of cortical tubules. B) Radii of viable glomeruli.

## Discussion

Image segmentation allows for the detection and classification of histologic structures and is considered a foundational step in the development of computational pathology and an absolute requirement for automated histomorphometric analysis. Generally, segmentation methods can be classified as either semantic segmentation or instance segmentation. In semantic segmentation, a classification label is assigned to every pixel in the image, but this method is limited in that it cannot distinctly recognize two same-class entities that are abutting or overlapping. In contrast, instance segmentation is the task of distinctly recognizing abutting/overlapping objects as unique entities. However, such algorithms are typically unable to model multiple classes. Numerous studies have shown the undeniable utility of deep neural networks for segmentation tasks in digital pathology datasets.^13-22^ Yet, most prior networks were constrained either for semantic segmentation or instance segmentation alone, unable to leverage the strengths of each method in combination. More recently, the development and maturation of panoptic architectures has led to the ability to segment both semantic and instance objects simultaneously, allowing for a comprehensive approach to histomorphometric image analysis. In this work, we demonstrate the feasibility of using a panoptic segmentation neural network-based pipeline to accurately quantify a variety of histomorphometric parameters from WSIs of reference kidney tissue sections. The high segmentation performance of our model allowed us to take the first steps in defining reference morphometrics for healthy human kidneys on over 3 million nephrons.

Although a large amount of histomorphometric data can be extracted using our automated DL-based image analysis pipeline, the measurements are only useful if they have some type of biological or clinical relevance. Thus, we subsequently used simple regression analysis to identify relationships between histomorphometric parameters of healthy kidneys to patient age, sex, and serum creatinine. Several histomorphometric parameters that reflect glomerular and tubular size and glomerular density significantly correlated to patient serum creatinine levels. Our observations are consistent with prior studies, which indicate that glomerular and tubular size tends to be directly correlated to serum creatinine levels, while glomerular density (as estimated by the number of glomeruli per renal cortical area) is typically inversely correlated to serum creatinine levels.

Our study has a few major limitations. First is the definition and use of reference kidneys. Depending on the stringency of defining the criteria for reference kidney samples, true reference kidney samples are difficult to obtain given the ethical considerations. One alternative would be using autopsy kidneys; however, these samples typically have prominent degradation/decomposition artifact, which would likely confound histomorphometric analysis. In our study, we used kidney parenchyma from tumor nephrectomy specimens distanced from the tumor foci and screened for minimal abnormalities by a pathologist. Such specimens are likely the most readily available, although the age of the patients is skewed to older individuals. Also, analysis of a tumor that could potentially result in a mass effect-related issue like obstruction could confound the data even after a pathologist has screened for normal-appearing sections. Another limitation is that we used large tissue sections from nephrectomy specimens that may not easily translate to equivalent histomorphometric values seen on biopsy. Typical biopsies have a much smaller surface area, which results in significantly lower numbers of glomeruli, tubules, and vessels, as well as a high proportion of transected structures at the edge of the biopsy. Whether reference kidney histomorphometric values need to be re-established in smaller biopsy specimens in order to be useful in the clinical setting needs to be evaluated. Furthermore, we used specimens from three different institutions, which likely resulted in an institutional-specific batch effect. Further evaluation to examine in detail the histomorphometric effects of processing tissue at different institutions must be determined. Finally, some important patient demographic data were not readily available to us, such as patient weight or BMI.

To our knowledge, our work is the most comprehensive study to tabulate large-scale reference renal morphometry features with clinical significance using a large, diverse, multinational, highly quality controlled cohort of renal tissue biopsy images. Ultimately, reference kidney histomorphometric values require examination of a large cohort from various populations, which will likely be achievable by upscaling our current strategy. Such data would allow eventual quantitative or statistical definitions for certain types of pathologic entities. For instance, tubular atrophy could one day be defined as tubules with radii below a certain statistical threshold. Similarly, defining glomerulomegaly may be more straightforward and diagnosis may be aided by automated morphometry.

## Supporting information

Augmentation Strategy, Tested Features, Supplementary Figures

## Disclosures

The authors have no conflicts of interest to disclose.

## Funding

Pinaki Sarder’s work is supported by NIH-NIDDK grant R01 DK114485, R01 DK131189, R21 DK128668, via the opportunity pool funding mechanism, namely via the glue grant mechanism of the NIH-NIDDK Kidney Precision Medicine Project (KPMP) consortium grant U2C DK114886, via the KPMP Kidney Mapping and Atlas Project (KMAP) U01 DK133090, NIH-OD Human Biomolecular Atlas Project (HuBMAP) consortium Integration, Visualization & Engagement (HIVE) project OT2 OD033753, NIH/NCI Coordinating and Data Management Center for Acquired Resistance to Therapy Network U24 CA274159, and faculty start-up funding from University of Florida.

## Acknowledgments

We would like to thank Ms. Jessica Kirwan for assisting with scientific editing of the manuscript and preparing it for submission.

## Author contributions

BG conceptualized and implemented the study design, coordinated data acquisition, provided expert annotations, performed all computational methods implementations and study interpretations, and wrote the first draft of the manuscript. KYJ co-designed the reference study, provided kidney reference samples and clinical metadata, conducted data quality control, expert annotation, interpretation of statistical findings, and provided clinical feedback on the overall work and manuscript. JZ and TOB assisted with design and interpretation of statistical analyses and reviewed and approved the final draft of the manuscript. SJ provided human kidney samples, assisted in reference study design, and reviewed and approved the final draft of the manuscript. SSH and LR provided kidney reference samples and clinical metadata and reviewed and approved the final draft of the manuscript. NL conducted segmentation tests, contributed to the manuscript, and reviewed and approved the final draft. MLW assisted with collection of human kidney samples and clinical metadata reviewed and approved the final draft of the manuscript. PS conceived the reference study, coordinated the study organization and data acquisition, provided feedback on the study design and manuscript preparation, and reviewed and approved the final draft of the manuscript.

## Notes

https://bit.ly/3YD4r6a

### Competing Interest Statement

The authors have declared no competing interest.

