## Supplementary material for "Correlating Deep Learning-Based Automated Reference Kidney Histomorphometry with Patient Demographics and Creatinine": Augmentation Strategy, Tested Features, Supplementary Figures

The augmentation procedure used to train the panoptic network, and referenced in *Training* of the main manuscript, is defined below in the order that the operations occurred to the image. First, to address variation in staining preparation, we added a random offset to the hue channel (of the HSV space) selected using a random normal distribution centered on zero with standard deviation 0.05. Then, to address brightness variation, we used gamma adjustment on the lightness channel (from the LAB colorspace) selected from a separate random normal distribution centered on one with standard deviation 0.025. Each patch had 90% probability of being subject to this procedure. Next, we used the ImgAug python package to add a number of general augmentation procedures, including elementwise noise, impulse noise, course dropout, Gaussian blurring, and sharpening[1]. The sequence of augmentations was as follows. Sometimes (50% probability), one of elementwise noise (values = (-15, 15), per_channel = 0.5), impulse noise (Pr = 0.05), or course dropout (Pr = 0.02, size_percent = 0.5) was selected randomly and uniformly and applied to the image. Then, sometimes (50% probability), one of Gaussian blur (sigma = (0.0, 3.0)) or sharpening (alpha = (0, 1), lightness = (0.75, 2.0)) was selected uniformly and randomly and applied. Finally, images were subjected to random horizontal flipping with 50% probability, which concluded the augmentation sequence.

### Tested Features

***Supp. Table 1.*** **Full list of tested morphometrics and reference values (N = 79 subjects, one whole slide image per subject).**

| | Institution 1 (n=43) | Institution 2 (n=8) | Institution 3 (n=28) | Combined (N=79) |
|---|---|---|---|---|
| ***Patient Characteristics*** | | | | |
| **Male sex** | 29 (67.44%) | 5 (62.50%) | 20 (71.43%) | 54 (68.35%) |
| **Age** | 59.16 ± 11.95 | 60.63 ± 13.32 | 53.68 ± 11.94 | 57.37 ± 12.25 |
| **SCr** | 1.07 ± 0.24 | 0.81 ± 0.28 [a] | 0.86 ± 0.14 [a] | 0.97 ± 0.24 |
| ***Glomerular histomorphometrics*** | | | | |
| **Glomeruli (per cortical mm$^2$)** | 2.59 ± 0.64 | 1.86 ± 0.57 [a] | 2.57 ± 0.80 [b] | 2.51 ± 0.72 |
| **Average glomerular area (µm$^2$)** | 18833 ± 3765 | 23836 ± 6113 [a] | 19248 ± 3361 [b] | 19487 ± 4134 |
| **Standard deviation glomerular area (µm$^2$)** | 9653 ± 3879 | 10863 ± 2699 | 8451 ± 1846 | 9350 ± 3242 |
| **Average glomerular radius (µm)** | 63.8 ± 6.81 | 72.08 ± 10.73 [a] | 65.93 ± 5.98 | 65.39 ± 7.33 |
| **Standard deviation glomerular radius (µm)** | 17.08 ± 2.63 | 19.62 ± 3.74 | 16.88 ± 2.94 | 17.27 ± 2.93 |
| **Sclerotic glomeruli (per cortical mm$^2$)** | 0.19 ± 0.14 | 0.08 ± 0.06 | 0.24 ± 0.21 | 0.20 ± 0.17 |
| **Average sclerotic glomerular area (µm$^2$)** | 8252 ± 1712 | 9222 ± 1883 | 7780 ± 1579 | 8188 ± 1713 |
| **Standard deviation sclerotic glomerular area (µm$^2$)** | 4159 ± 1325 | 5293 ± 3085 | 2797 ± 1319 [a,b] | 3804 ± 1751 |
| **Average sclerotic glomerular radius (µm)** | 42.04 ± 4.66 | 44.69 ± 5.13 | 42.62 ± 4.68 | 42.51 ± 4.72 |
| **Standard deviation Sclerotic glomerular radius (µm)** | 11.56 ± 2.03 | 13.03 ± 3.45 | 8.33 ± 3.89 [a,b] | 10.59 ± 3.37 |
| **Glomerulosclerosis ratio** | 0.07 ± 0.06 | 0.04 ± 0.04 | 0.08 ± 0.07 | 0.07 ± 0.06 |
| **Glomerular proportion of cortex** | 0.05 ± 0.01 | 0.04 ± 0.01 | 0.05 ± 0.01 | 0.05 ± 0.01 |
| ***Tubulointerstitial morphometrics*** | | | | |
| **Tubules (per cortical mm$^2$)** | 173.53 ± 34.40 | 132.92 ± 38.17 [a] | 189.58 ± 50.16 [b] | 175.10 ± 43.57 |
| **Average cortical tubular area (µm$^2$)** | 3269 ± 612 | 4219 ± 1141 [a] | 2476 ± 722 [a,b] | 3084 ± 885 |
| **Standard deviation cortical tubular area (µm$^2$)** | 3495 ± 1002 | 5752 ± 3652 [a] | 2809 ± 1232 [b] | 3481 ± 1719 |
| **Average cortical tubular radius (µm)** | 20.95 ± 1.53 | 23.43 ± 2.39 [a] | 17.92 ± 2.31 [a,b] | 20.13 ± 2.62 |
| **Standard deviation cortical tubular radius (µm)** | 7.65 ± 1.02 | 8.93 ± 1.67 [a] | 7.00 ± 1.38 [b] | 7.55 ± 1.33 |
| **Average medullary tubular area (µm$^2$)** | 1768 ± 506 | 2449 ± 453 [a] | 1352 ± 464 [a,b] | 1689 ± 579 |
| **Standard deviation medullary tubular area (µm$^2$)** | 2098 ± 1027 | 3403 ± 1095 [a] | 1742 ± 1084[,b] | 2104 ± 1141 |
| **Average medullary tubular radius (µm)** | 15.37 ± 1.50 | 17.75 ± 1.48 [a] | 12.69 ± 1.20 [a,b] | 14.66 ± 2.13 |
| **Standard deviation medullary tubular radius (µm)** | 5.14 ± 1.01 | 6.32 ± 0.54 [a] | 4.68 ± 0.88 [b] | 5.10 ± 1.03 |
| **Tubular proportion of cortex** | 0.55 ± 0.05 | 0.52 ± 0.07 | 0.45 ± 0.09 [a,b] | 0.51 ± 0.08 |
| **Average lumen to wall ratio** | 0.27 ± 0.05 | 0.37 ± 0.05 [a] | 0.23 ± 0.04 [a,b] | 0.27 ± 0.06 |
| **Tubular proportion of medulla** | 0.49 ± 0.10 | 0.49 ± 0.08 | 0.38 ± 0.07 [a,b] | 0.45 ± 0.10 |
| **Interstitial proportion of cortex** | 0.16 ± 0.10 | -0.45 ± 0.13 [a] | 0.31 ± 0.06 [a,b] | 0.15 ± 0.23 |
| **Nephron count** | 202.79 ± 39.70 | 149.92 ± 29.42 [a] | 218.45 ± 50.52[b] | 202.99 ± 46.74 |
| **Interstitial proportion of medulla** | 0.51 ± 0.10 | 0.51 ± 0.08 | 0.62 ± 0.07 [a,b] | 0.55 ± 0.10 |
| ***Vascular morphometrics*** | | | | |
| **Arteries(ioles) per cortical mm$^2$** | 5.96 ± 1.59 | 4.04 ± 1.14 [a] | 6.54 ± 1.98 [b] | 5.97 ± 1.82 |
| **Arter(iole) proportion of cortex** | 0.04 ± 0.02 | 0.03 ± 0.01 | 0.04 ± 0.02 | 0.04 ± 0.02 |
| **Arteriole proportion of medulla** | 0.03 ± 0.01 | 0.03 ± 0.02 | 0.01 ± 0.01 [a,b] | 0.02 ± 0.01 |
| **Cortical glomerulus to tubule ratio** | 0.09 ± 0.02 | 0.08 ± 0.02 | 0.11 ± 0.02 [a,b] | 0.09 ± 0.02 |

Data are represented as mean ± standard deviation, if not indicated otherwise. *Standard deviation for the population of glomeruli within a single case. [a] Adjusted p-value ≤ 0.05 compared to Institution 1. [b] Adjusted p-value ≤ 0.05 compared to Institution 2.

### Supplementary Figures

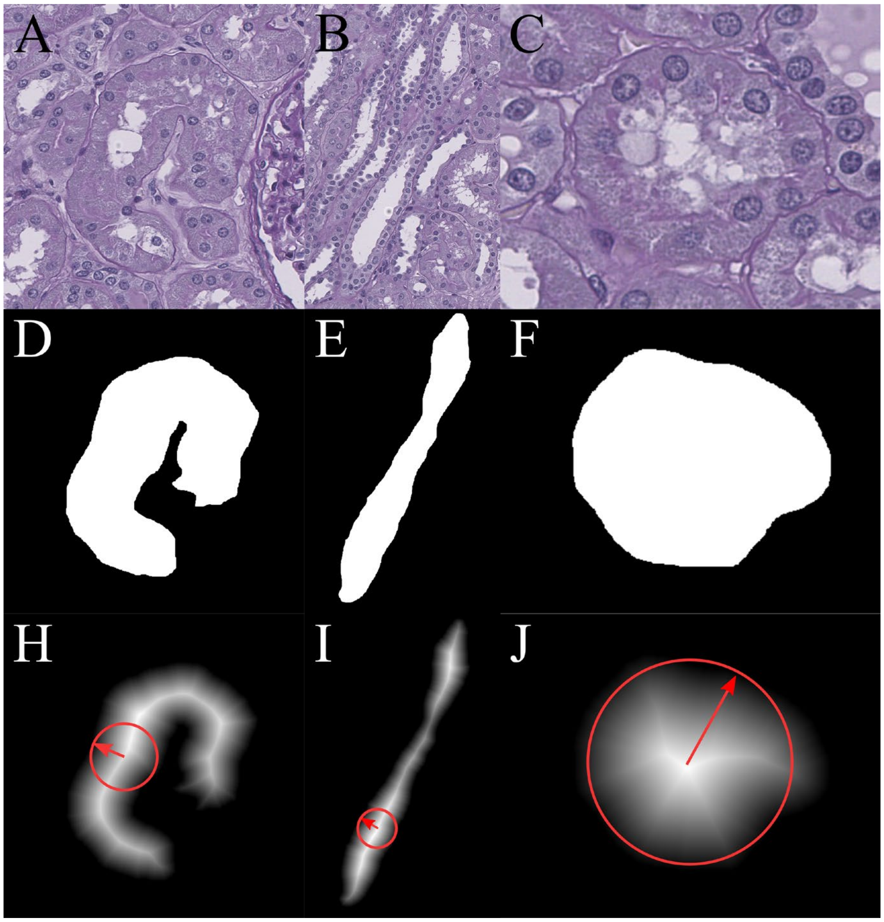

***Supp. Fig. 1.*** **Examples of distance transform applied to various tubular segments.** A-C) Original histology images showing a curled, straight, and cross-sectioned tubule, respectively. D-F) Binary segmentation masks of the tubules in top row. H-J) Distance transformation output, an intensity image with each pixel valued as the distance to closest boundary point. Red arrow and circle annotations correspond to the approximate maximum radius and its circular inscription for each tubule.

#### Distance transform exemplars

In this work, we chose the distance transform to measure object radius/diameter. ***Supp. Figs. 1A-1C*** show three examples of tubules, one curled unto itself, one straight, and one directly cross-sectioned. The corresponding segmentation masks of these tubules are provided in ***Figs. 1D-1F.*** The maximum value of the distance transformation identifies the maximum radius of a circle that could be inscribed in the object, as shown in ***Figs. 1H-1J***. This quantification allows an accurate measurement of the tubular diameter/radius regardless of the angle at which the tubule was sectioned.

### Segmentation of other histological stains

We included a limited amount of histological staining other than PAS in the training of our segmentation network. We provide a brief qualitative demonstration of network performance on hematoxylin and eosin (H&E), trichrome, and silver stained kidney with very low training data.

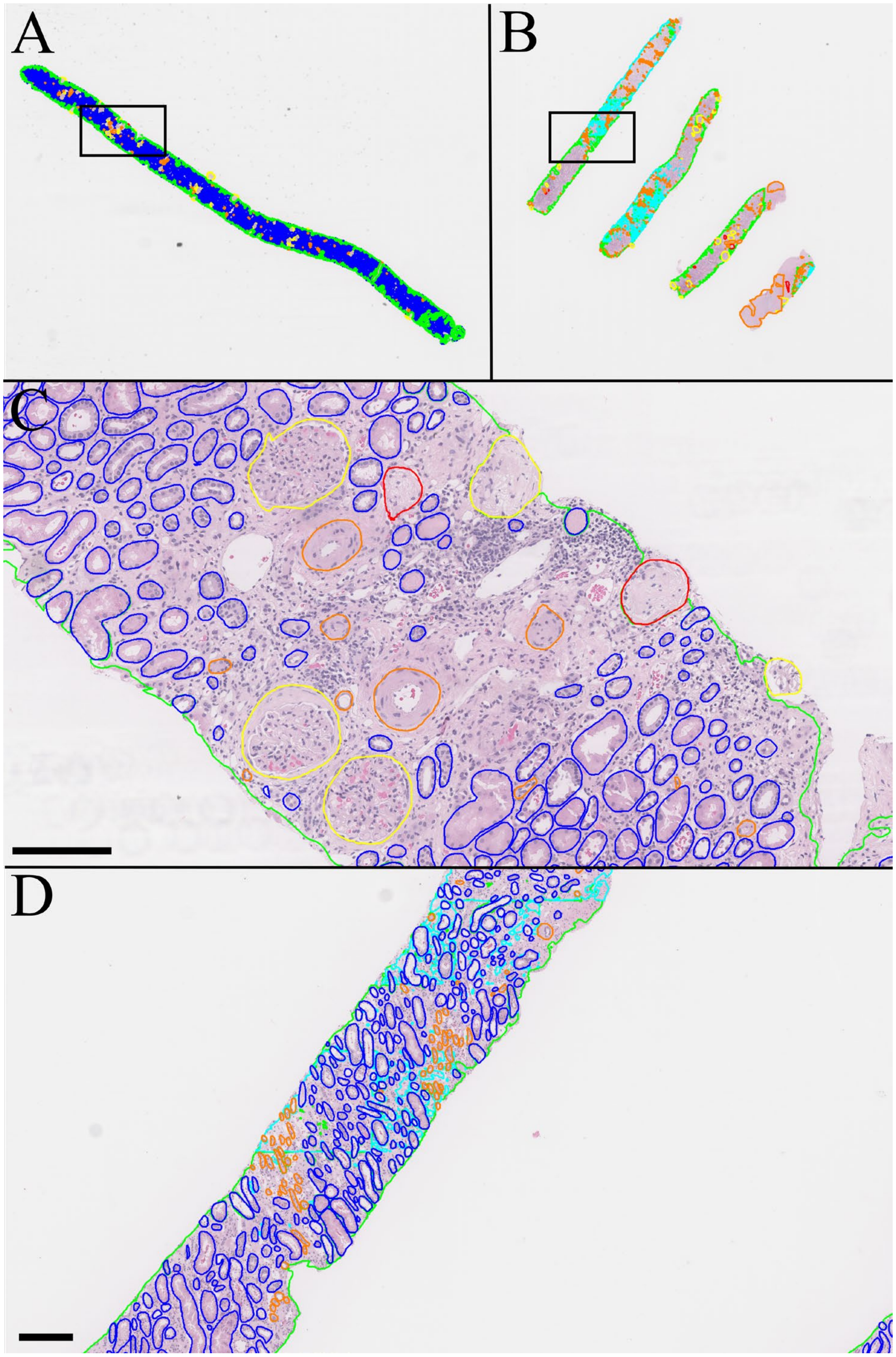

***Supp. Fig. 2.*** **Holdout segmentation of H&E biopsy after inclusion of 23 small training regions totaling 34mm$^2$.** A) Thumbnail image of all segmentation boundaries for a biopsy with chronic kidney disease (CKD). B) Thumbnail segmentation of a second biopsy with medulla. C) 12X magnification zoomed inset for the rectangular region in A. D) 4x magnification zoomed inset for the rectangular region in B. Green: cortical interstitium, cyan: medullary interstitium, yellow: viable glomerulus, red: sclerotic glomerulus, blue: tubule, orange: artery/arteriole. Scale bars 150µm.

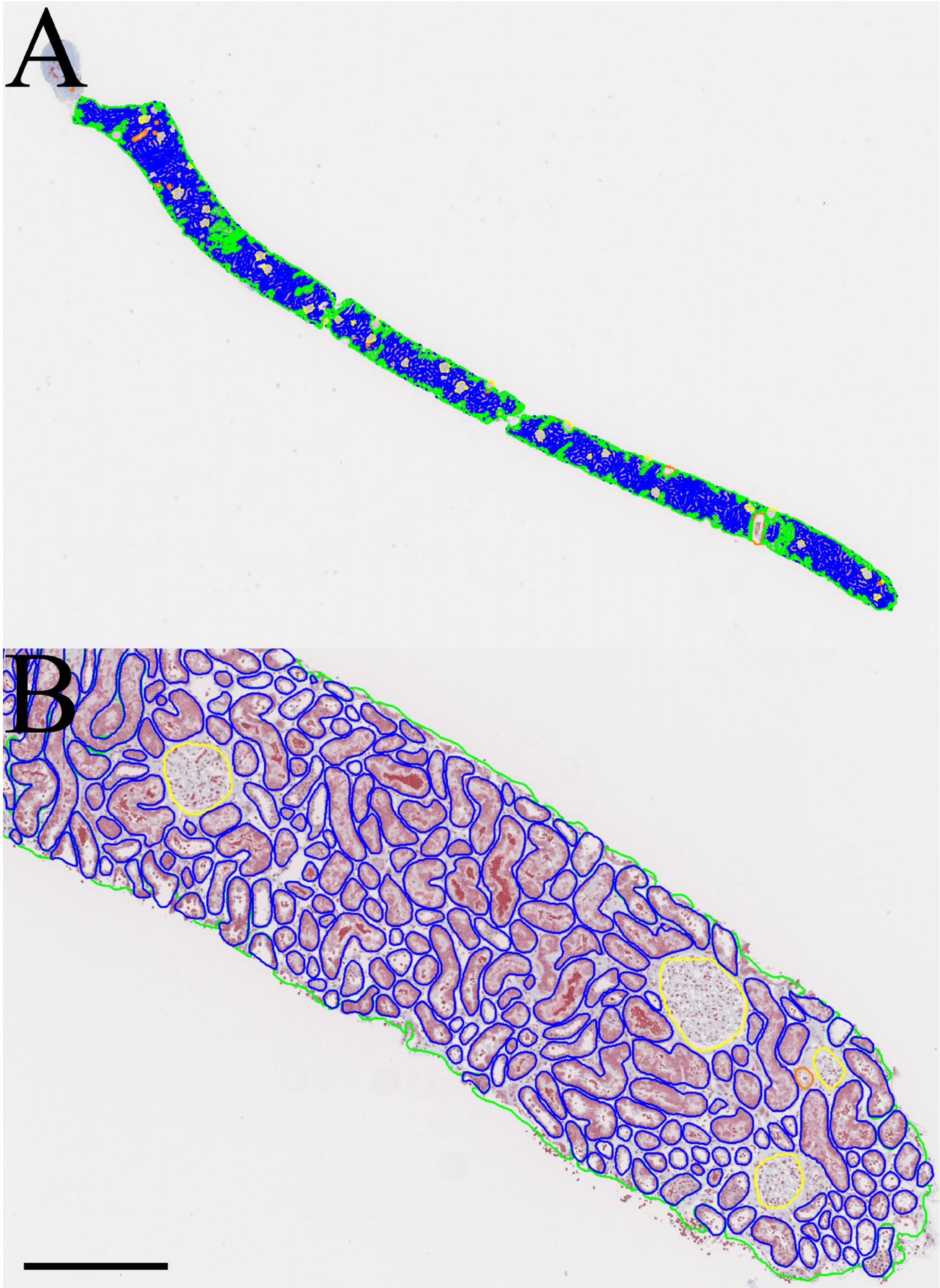

***Supp. Fig. 3.*** **Holdout segmentation of trichrome biopsy after inclusion of four annotated trichrome training patches totaling only 1.4mm$^2$ of tissue.** A) Thumbnail image of whole slide predictions. B) Zoomed inset showing individual segmentation boundaries. Green: cortical interstitium, yellow: viable glomerulus, blue: tubule, orange: artery/arteriole. Scale bar 300µm.

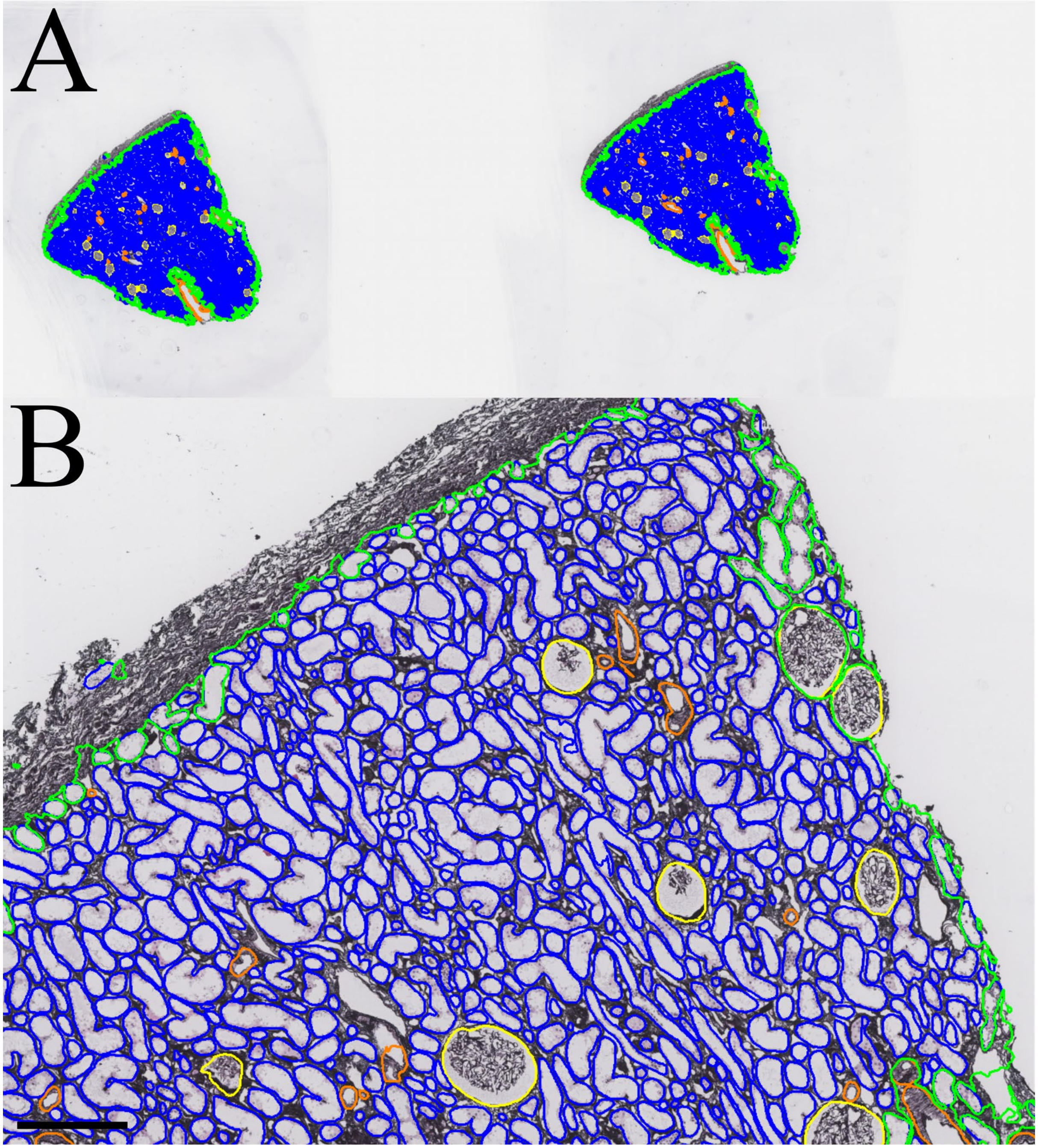

***Supp. Fig. 4.* Holdout segmentation of silver biopsy after inclusion of two annotated silver training patches totaling only 2mm$^2$ of tissue.** A) Thumbnail image of whole slide predictions. B) Zoomed inset showing individual segmentation boundaries. Green: cortical interstitium, yellow: viable glomerulus, blue: tubule, orange: artery/arteriole. Scale bar 250µm.
